# Optimizing Bispecific Antibody Expression via Multi-Omics Analysis and Vector Redesign

**DOI:** 10.1101/2025.11.10.687454

**Authors:** Jeremy J. Gam, Michelle M. Chang, Dinghai Zheng, Jennitte Stevens, Alec A.K. Nielsen, Kevin D. Smith

## Abstract

Bispecific antibodies are a growing class of therapeutics that simultaneously engage two targets. However, their complex molecular structures pose challenges for production in Chinese hamster ovary cells, the current industry standard for biologics manufacturing. Here we present a case study of three IgG-scFv format bsAbs expressed in CHO cells, in which one candidate exhibited markedly lower titers despite high sequence homology to the other two. Using multi-omics analysis (RNA sequencing, splicing prediction, codon optimization assessment, and motif screening) to investigate potential causes, we identified several likely mechanisms for poor expression, including aberrant splicing motifs, ribosome pausing sites, and suboptimal codon usage. Through targeted protein and DNA sequence engineering, we generated a revised variant with an 11-fold increase in stable expression titers. This work demonstrates that integrating sequence-level bioinformatic and synthetic biology diagnostics can directly improve manufacturability, providing a generalizable framework for resolving hidden expression liabilities in complex biologics.

## Introduction

### Immunotherapy has revolutionized medicine, and monoclonal antibodies have long been a driving force

More than 100 monoclonal antibodies, or mAbs, have been approved in the United States for treating diseases ranging from viral infections to cancer and inflammation. [1] mAbs function by selectively binding a single target epitope, which either sterically blocks access to the target or recruits the immune system for a downstream response. A single-target approach can be problematic for diseases such as cancer and autoimmune disorders, however, because the targeted epitope can mutate to evade or resist the mAb.

Bispecific antibodies, or bsAbs, have built upon the success of conventional monoclonal antibodies while minimizing some of their shortcomings. In contrast to mAbs, bsAbs bind two targets simultaneously. In a process known as T-cell engagement, for example, bsAbs simultaneously bind surface epitopes on both T-cells and diseased cells, thereby redirecting potent T-cell cytotoxicity towards target cells via induced proximity. Alternatively, with immune-checkpoint modulation, simultaneous binding of two immune checkpoint regulators (e.g. PD-1 or CTLA4) can synergize the response, resulting in a reduced ability for tumor cells to evade T-cells and the immune system. [2, 3] As of 2025, more than 14 bsAbs have garnered FDA approval and over 500 more are in clinical trials - most of which are designed to treat cancer. [4, 5]

Chinese hamster ovary (CHO) cells have become the standard expression system for bsAbs due to their robust protein expression capabilities, human-like protein folding and post-translational modifications, and existing infrastructure for commercial-scale production. [6, 7] Engineered CHO cell lines also have established genetic backgrounds and auxotrophies, and there is a long track record of using these cells in combination with genetic tools and bioreactor process optimization. [8, 9, 10] Despite the adoption of CHO for general biologics manufacturing, it can still be difficult to make bsAbs at high yields and qualities. [11, 12]

bsAbs have complex structures and do not always fold or assemble correctly inside the cell. [12] Most bsAbs are designed as two or more protein chains that must be expressed at the proper ratio and then complexed to form the correct quaternary structure. Improper folding within individual chains, chain expression imbalance, incomplete assembly, or mispairing of chains can result in reduced yield of functional bsAb molecules. [13] Increased molecular complexity also increases the likelihood for proteolytic clipping, aggregation, and product-related impurities, which necessitates more stringent purification and quality control processes to ensure drug efficacy and safety. [14, 15]

Numerous protein engineering strategies have been developed to reduce bsAb complexity or enforce proper pairing of the protein chains in order to increase yield. bsAbs can be designed with fewer chains by fusing chains together with flexible linkers (e.g., scFv fusions to heavy chains). [16, 17] Proper pairing of chains can also be enforced by introducing a knob-in-hole structure within the Fc by engineering single sterically matched mutations in the CH3 domain, by incorporating charge mutations in the CH3 domain, or by swapping the CH1 and CL domains of the Fab (e.g., in the CrossMAb architecture). [18, 19, 20] bsAb titers can also be increased independently of protein engineering by optimizing gene expression cassettes to properly balance the chain expression ratios, increasing the copy number or the expression per copy of the integrated gene expression cassettes, or by optimizing bioreactor production processes. [21, 22, 23] It is of course important to avoid negatively impacting binding specificity or product quality in pursuit of higher titers, so in addition to protein engineering to improve the properties of bsAbs, understanding mechanisms that lead to poor rates of protein translation, folding, and secretion is another important consideration to help improve the manufacturability of these important class of therapeutics.

Here we present a case study highlighting a group of three bsAbs with similar architectures, one difficult to express and the other two more easily expressed. We then used computational and genetic engineering techniques to determine potential biological mechanisms underpinning the production difficulties. Lastly, using these insights, we generated a revised bsAb sequence with 11-fold greater titers than the initial design. Our findings demonstrate that a combination of protein engineering to eliminate specific problematic motifs and sequence optimization can result in additive improvements in bsAb productivity.

## Materials and Methods

### Cell Culture

Proprietary CHO GS knockout host cells were obtained from Amgen and cultured in proprietary Growth Media containing L-glutamine. The suspension cultures were maintained in a humidity control shaker incubator (New Brunswick s41i) at 36°C, 5% CO_2_, and 130 rpm.

### Plasmid construction

An expression backbone containing a GS expression cassette, and two hEF1a transcription units was assembled using a combination of BsaI Golden Gate assembly (NEB Cat. No. R3733S and Promega Cat. No. M1794) followed by Gibson assembly (NEB Cat. No. E2621L). Codon optimized sequences were codon optimized using Genscript’s codon optimization algorithm. bsAb HC and LC sequences were synthesized into precursor plasmids and assembled into the expression backbone using BbsI Golden Gate assembly (NEB Cat. No. R0539S). DNA assembly reactions were transformed into NEB 10-beta Electrocompetent E. coli (Cat. No. C3020K). Insertion and identity of bsAb HC and LC sequences was confirmed by Sanger sequencing. Transfection-grade DNA was prepared using a QIAGEN Plasmid Plus Midi Kit (Cat. No. 12945).

### Transfection and titer measurement

Stable pools were generated by random integration. Non-linearized plasmids (4 ug) were transfected into CHO host cells (4e6 cells) using Lipofectamine LTX (8 uL, Gibco Cat. No. 15338100) in Opti-MEM I Reduced Serum Medium (2 mL, Gibco Cat. No. 31985070) in a non-treated 6-well plate and incubated at 36°C, 5% CO_2_, and 130 rpm. Transfected cells were supplemented with Growth Media (2 mL) after 6 hours. Cells were placed into proprietary Selection Media without glutamine 48-72 hours post transfection and split twice weekly until recovered. Stable pools were then moved to Growth Media for batch production.

For transient production, CHO host cells were transfected using the same method as in the stable pool generation. Cell-free supernatants were titered 72 hours post transfection.

Titer measurement of the bsAbs was carried out on Octet Red96e and Octet RH96 Bio-Layer Interferometry instruments using Octet ProA biosensors (Sartorius). Cell-free supernatants were used from harvested transient or stable production cultures.

### Batch and fed-batch production

For batch production, the cultures were seeded at 1.8e6 cells/mL in Growth Media and supplemented with 5% (v/v) Feed Media and 8 g/L glucose and harvested on days 6-8. For a 14-day fed batch production process, the cultures were seeded at 1e6 cells/mL at 65% of the desired final culture volume in Selection Media. Cultures were supplemented with 7% (v/v) Feed Media (final volume) and 8 g/L glucose (current volume) on days 3, 6, 8, and 10. Cultures were supplemented with 3% (v/v) Feed Media (final volume) and 6 g/L glucose (current volume) on days 11, 12, and 13.

### RNA-seq and analysis

Cell pellets were flash frozen in liquid nitrogen. RNA was extracted and analyzed on the Illumina HiSeq system through the Standard RNA-Seq package provided by Genewiz.

### DNA sequence analysis and predictions

DNA sequence analyses and predictions were performed using a suite of bioinformatic tools and published metrics. These included assessments of codon adaptation index [24], translational pausing identification by simple text search, mRNA secondary structure [25], splice site strength, and RIDD motifs [26], as described in previously published studies.

**Figure 1:**
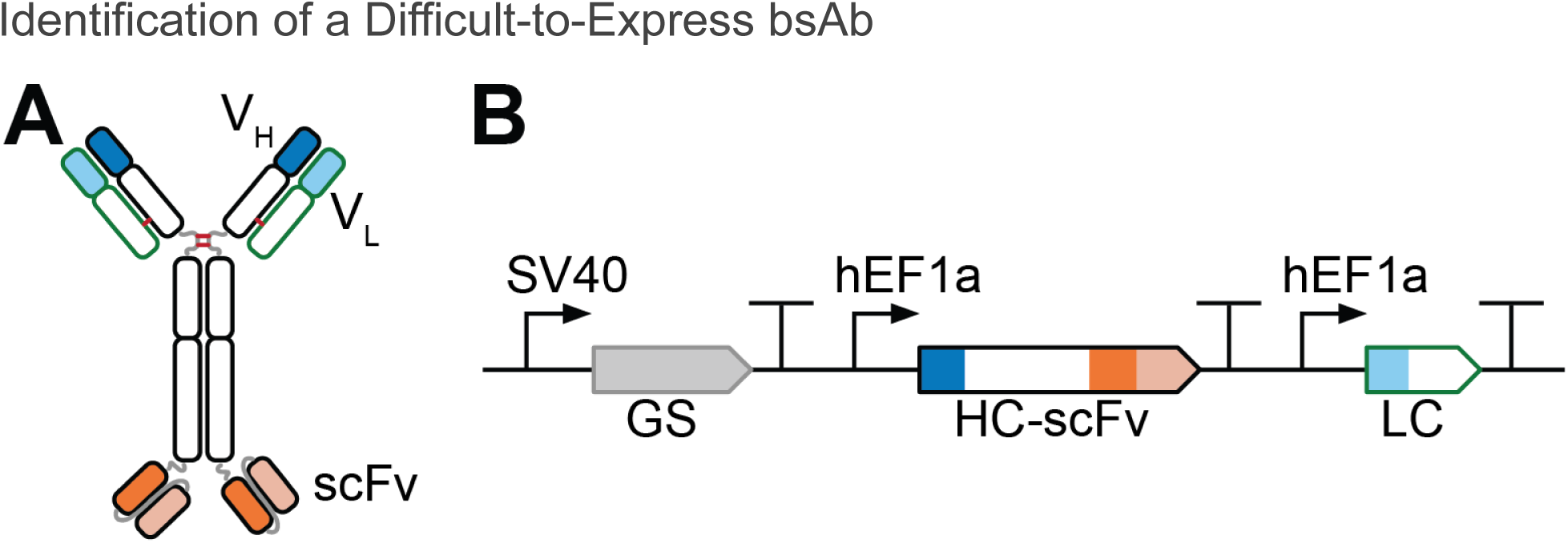
bsAb architecture and expression plasmid design A) IgG-scFv bsAb architecture. The light chain (green outlines) is a typical mAb-like design, whereas the engineered heavy chain (black outlines) contains an scFv fused to the C-terminus of a typical mAb-like heavy chain via a flexible linker. Variable regions map to the typical mAb-like variable region (blue/cyan fill) and the scFv (orange/light orange fill). V_H_ domains are dark blue and orange, whereas V_L_ domains are cyan and light orange. Inter-chain disulfide bonds are illustrated as red lines. B) Plasmid schematic for the bispecific antibody expression constructs. GS denotes the glutamine synthetase selection marker. Coloring of regions and outlines are the same as in (A).

## Results

### Identification of a Difficult-to-Express bsAb

Dozens of different bsAb architectures have been developed to date, each with unique target affinities, pharmacokinetics, and effector functions. [16, 17, 27] Here we focus on three bsAbs of the IgG-scFv design (commonly referred to as the Morrison format), which is based on a monoclonal antibody with a single-chain variable fragment (scFv) fused to the C-terminus of the heavy chain via a flexible (G4S)2 linker (Figure 1). [28] These three bsAbs all have shared regions of high amino acid sequence homology, with the light chains and IgG domains being identical between the three molecules, and with differences limited to specific stretches within the Complementarity-Determining Regions (CDRs) of the Vh, scFv Vh, and scFv Vl regions.

Plasmid vectors for the three bsAbs (bsAb-A, bsAb-B, and bsAb-C) were constructed. Each vector contained expression cassettes for heavy chain (HC) and light chain (LC) with either native codons or codons optimized for CHO expression using an optimization algorithm from a leading DNA synthesis vendor. Each vector was randomly integrated into a proprietary CHO-GS knockout cell line. Then, cells were selected in glutamine-deficient media, followed by 8-day batch production culture (Figure 2). Titers were measured by the Octet system. bsAb-A achieved the highest titers even with the original coding sequences, whereas bsAb-B required codon optimization to achieve a 260% increase in titer to reach similar titers to bsAb-A. bsAb-C resulted in the lowest titer, and there was no significant improvement with codon optimization, achieving only 13% of the titer observed with the codon optimized versions of bsAb-A and bsAb-B.

**Figure 2:**
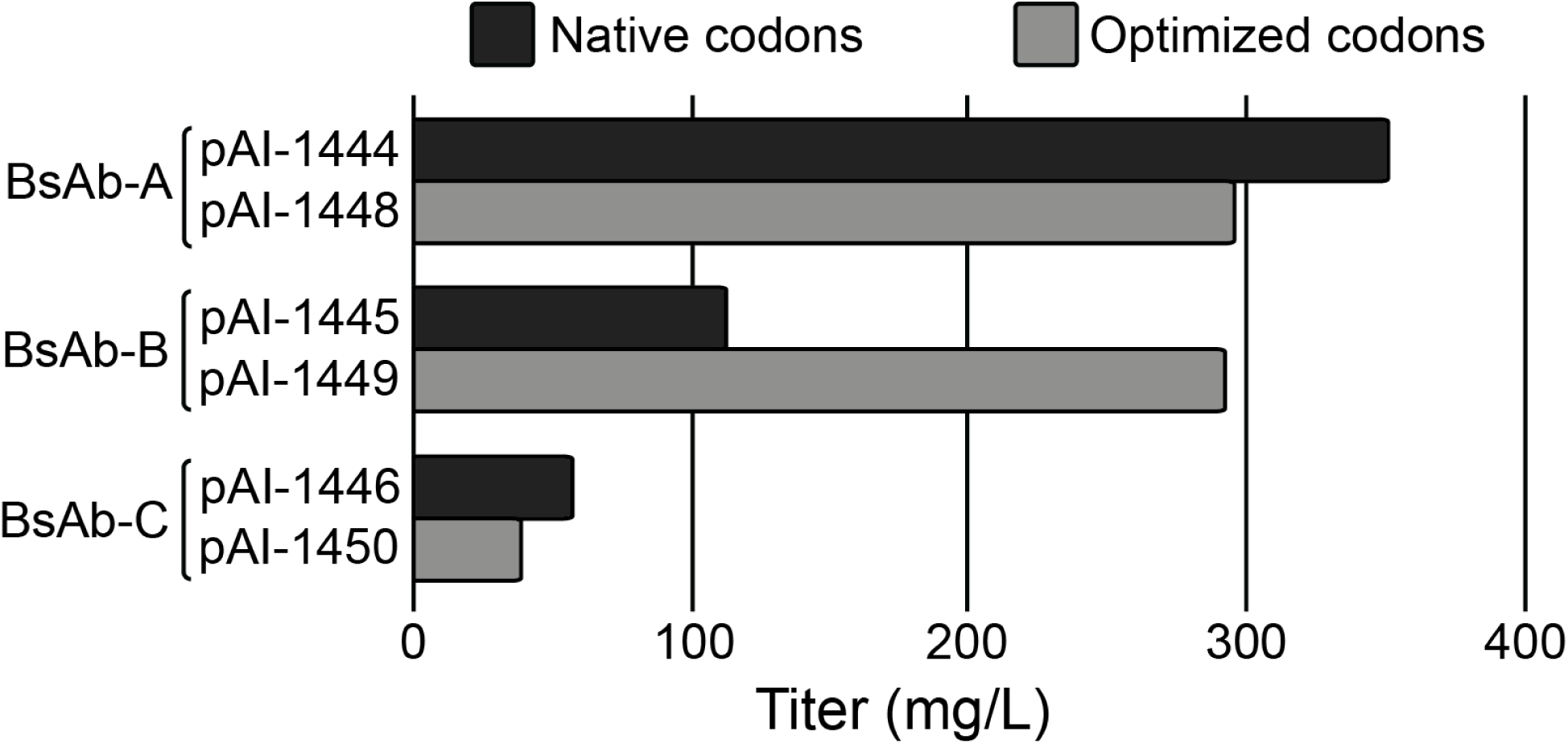
Stable pool titers from 8 day batch cultures. CHO lines were randomly integrated with plasmids containing the original nucleotide sequences (pAI-1444 to pAI-1446) or plasmids with codon optimized nucleotide sequences (pAI-1448 to pAI-1450).

### Bioinformatic-based Hypothesis Testing

We systematically evaluated the bsAb-C sequence using bioinformatic tools to generate a list of plausible hypotheses that could explain its low titers in batch cultures (Table 1). We used RNA-seq to investigate differential gene expression for each of the bsAbs, relative to the host CHO line. Additionally, we searched for differences in regulation of GSEA hallmark gene sets and KEGG pathways for each of the pools.

**Table 1:**
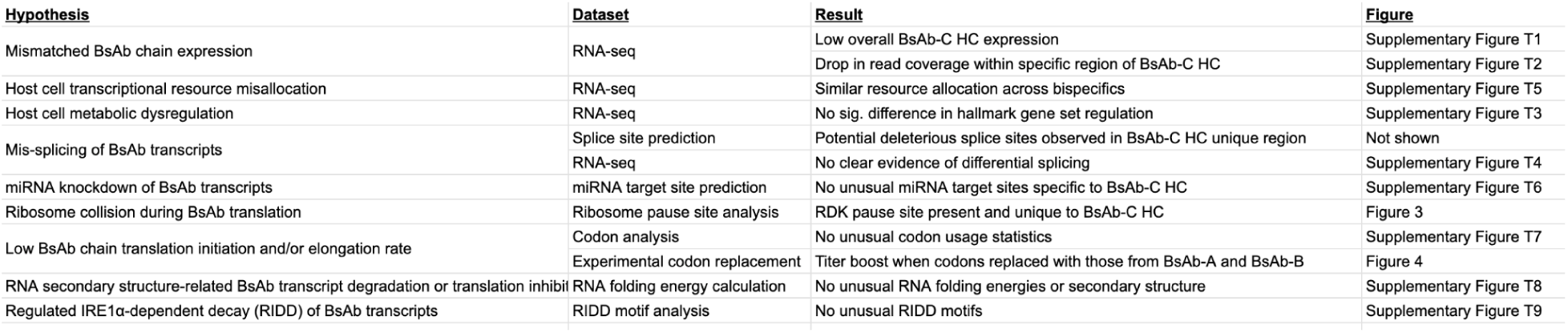
Summary of hypotheses for low bsAb-C titers explored bioinformatically.

Our analyses showed no prominent differences in host gene expression or gene sets for bsAb-C compared to the other bsAbs (Supplementary Figure S1). Expression of LC, measured as transcript per million (TPM), appeared consistent across the three bsAbs. However, we observed a 2-fold reduction in transcript abundance for bsAb-C HC compared to other bsAb HC transcripts (Supplementary Figure S2). Furthermore, we observed a significant dip in RNA-seq read coverage across a portion of bsAb-C HC, which was not observed in the other bsAb HC transcripts (Supplementary Figures S3-4).

We analyzed RNA splicing patterns within bsAb-C HC to identify potential deleterious splicing that could result in a reduction in RNA-seq read coverage and a truncated coding sequence, which may explain the low observed titer. SpliceRover, a splicing prediction algorithm, identified several splicing isoforms within the 3’ end of bsAb-C HC that could be detrimental, and which were unique to bsAb-C. [29] However, there was no clear evidence within the RNA-seq dataset of differential splicing that might explain the dip in RNA-seq read coverage. Nonproductive splicing isoform(s) may be unstable, however, resulting in low numbers of junction reads flanking the dip in bsAb-C HC. To experimentally test the possible effects of mis-splicing, an expression plasmid (pAI-2649) was generated with mutations to disrupt the five predicted splicing motifs.

Ribosome pausing and ensuing collisions can lead to decay of both mRNA and nascent peptides via no-go decay and the ribosome-associated quality control pathway. [30, 31, 32] Ribosome collisions result in disomes – which are two ribosomes stacked together due to translational pauses – and sequencing of disome-associated RNA has revealed polypeptide motifs associated with ribosome pausing, collisions and decay. Motifs fall into two classes: P-P/G/D and R-X-K. [33] We identified many instances of P-P/G/D motifs across all bsAbs that we tested, but only bsAb-C HC contained a R-X-K motif, specifically R-D-K at codon positions 552-554 (Figure 3). We hypothesized that the R-D-K motif may contribute to no-go decay and result in a decrease in both mRNA and protein abundance.

Sub-optimal codon choice could also explain the low titer for bsAb-C. As a first step to understanding codon usage, we compared HC amino acid sequences for bsAb-C to bsAb-A and bsAb-B. bsAb-C showed high homology to bsAb-B from the N-terminus to the (G4S)2 linker in the middle of the scFv, and high homology to bsAb-A from the (G4S)2 linker to the C-terminus (Figure 4A). Overall, the bsAb-C HC amino acid sequence only differed from bsAb-A or bsAb-B by a total of 32 amino acids, all localized within four distinct regions (6 to 49 residues in length) corresponding to specific stretches within the CDRs of the Vh, scFv Vh, and scFv Vl domains. We hypothesized that these four regions that were unique to bsAb-C could contain problematic amino acid sequences and/or coding sequences.

We explored several other hypotheses for low bsAb-C titers including unintended miRNA target sites, sub-optimal RNA secondary structure, and regulated IRE1-dependent decay of mRNA (RIDD). However, we determined that there are no miRNAs predicted to regulate bsAb-C strongly and specifically, RNA folding energies and local secondary structure of bsAb-C appears similar to the other bsAbs, and there are no unusual RIDD motifs present in bsAb-C (Supplementary Figures S5-8).

We also hypothesized that the design of the expression constructs themselves could impact titer. Transcription units positioned downstream from other transcription units can have reduced expression. [34] In the original design, the bsAb-C LC transcription unit was positioned downstream from the bsAb-C HC transcription unit. This could result in an improper HC and LC balance, which may exacerbate molecular assembly problems with bsAb-C. Therefore, we generated new constructs with the positions of bsAb-C HC and bsAb-C LC swapped to test if changing the HC/LC ratio improved expression.

In summary, our bioinformatic analysis resulted in prioritization of mis-splicing, codon usage, the order of HC and LC transcription units, and ribosome collision for further experimental testing. Revised expression plasmids were generated to test the potential impact of each.

**Figure 3:**
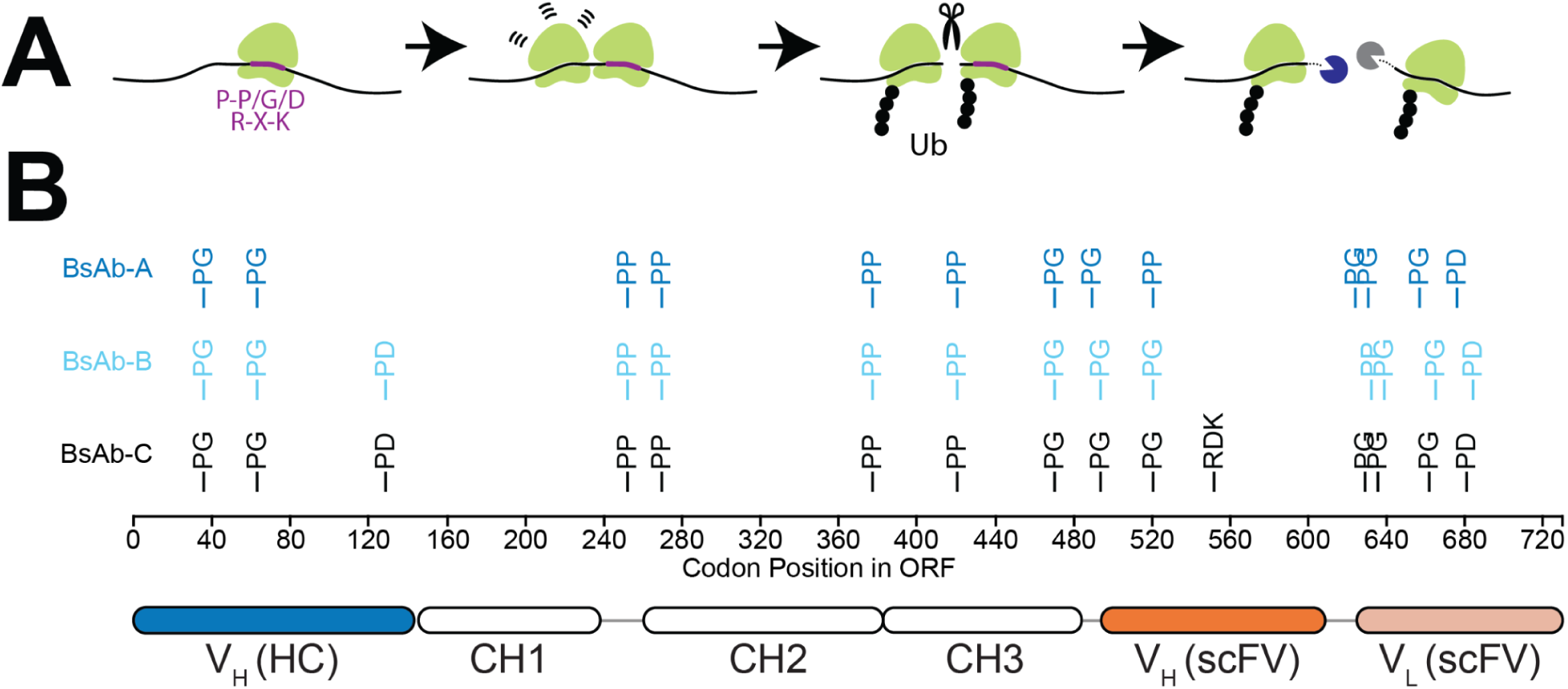
Predicted ribosome pause sites within heavy chain sequences A) Illustration of ribosome pausing and no-go decay. Pausing is enriched at certain sites, including stop codons and 3’ UTRs, polylysine tracts, P-P/G/D amino acid motifs, and R-X-K amino acid motifs. Pausing leads to ribosomal collisions and formation of disomes, ubiquitination of ribosome subunits, degradation of nascent peptides, RNA cleavage, RNA degradation, and ribosome recycling. B) P-P/G/D and R-X-K sites within the bsAb HC coding sequences. Amino acid motifs for ribosome pausing were identified in each HC chain. bsAb-B HC and bsAb-C HC contain an additional PD motif at positions 129-130 and bsAb-C HC contains the only R-X-K within the group at positions 552-554. Locations of bsAb variable and constant domains are indicated as ovals labelled V and CH for reference

### Sequence Determinants and Recovery of Low bsAb-C Bispecific Titers

Amino acid and DNA sequence alignments of bsAb-C to the other higher titer producing bsAbs confirmed the high sequence homology between bsAb-C HC and bsAb-B HC (N-terminus) and between bsAb-C HC and bsAb-A HC (C-terminus). They also revealed the amino acid sequence identity between bsAb-C LC and bsAb-B LC (Figure 4A).

Due to the high similarity between the aforementioned chains and the higher titers observed for bsAb-A and bsAb-B, we designed a chimeric sequence that replaces non-unique regions within bsAb-C with codon sequences from bsAb-A and bsAb-B, resulting in a construct with amino acid sequence identical to bsAb-C (Figure 4B, pAI-2636). Additionally, we created chimeras that replace one or more unique bsAb-C regions with those from bsAb-A and bsAb-B, resulting in constructs with very similar, but not identical, amino acid sequence compared to bsAb-C (Figure 4B, pAI-2637 through pAI-2647). If one or more of the four unique bsAb-C HC sequence stretches was detrimental to expression, we reasoned that the bsAb-A-B chimera (pAI-2639) should exhibit increased titer while chimeras incorporating bsAb-C-unique regions would exhibit low titer.

Surprisingly, the bsAb-A-B chimera exhibited low titer during transient production, suggesting that amino acid changes introduced into the scFv region reduced expression (Figure 4C, pAI-2639). Reintroducing the bsAb-C sequence, especially stretches 1 and 2, into the chimeras improved titers by greater than 500% (pAI-2636, pAI-2638, pAI-2646, pAI-2647). Therefore,we conclude that it is not simply the composition of amino acids in the scFv region of bsAb-C, rather we hypothesize that the codon usage within bsAb-C is having some impact on expression. We decided to evaluate codon usage as a parameter to test in follow-up stable production experiments.

Mutating the putative RDK ribosome pause site (within pAI-2636) to RDN (pAI-2648) resulted in negligible changes in titer during transient production. However, although both constructs showed similar Octet titers, they were assayed independently so we concluded that the detrimental effect of the RDK sequence could not be completely ruled out and should be tested further in a stable production format.

To evaluate the two remaining hypotheses from above, focusing on splicing and transcriptional unit positioning, we tested two additional expression plasmids. The pAI-2649 DNA sequence is similar to pAI-1450 (original bsAb-C HC) but with the RDK to RDN change and also several synonymous point mutations to eliminate splice motifs. Elimination of splice motifs showed little benefit to bsAb-C production. The pAI-2651 sequence is identical to pAI-2636, except that the locations of the LC and HC transcription units are swapped. We observed lower titers with pAI-2651, in which the HC was downstream of the LC, suggesting that swapping chains could have imbalanced the HC to LC ratio, potentially resulting in HC being too high and LC being too low. Consequently, we concluded not to pursue splicing or LC/HC transcriptional unit positioning in the follow-up stable production experiment.

**Figure 4:**
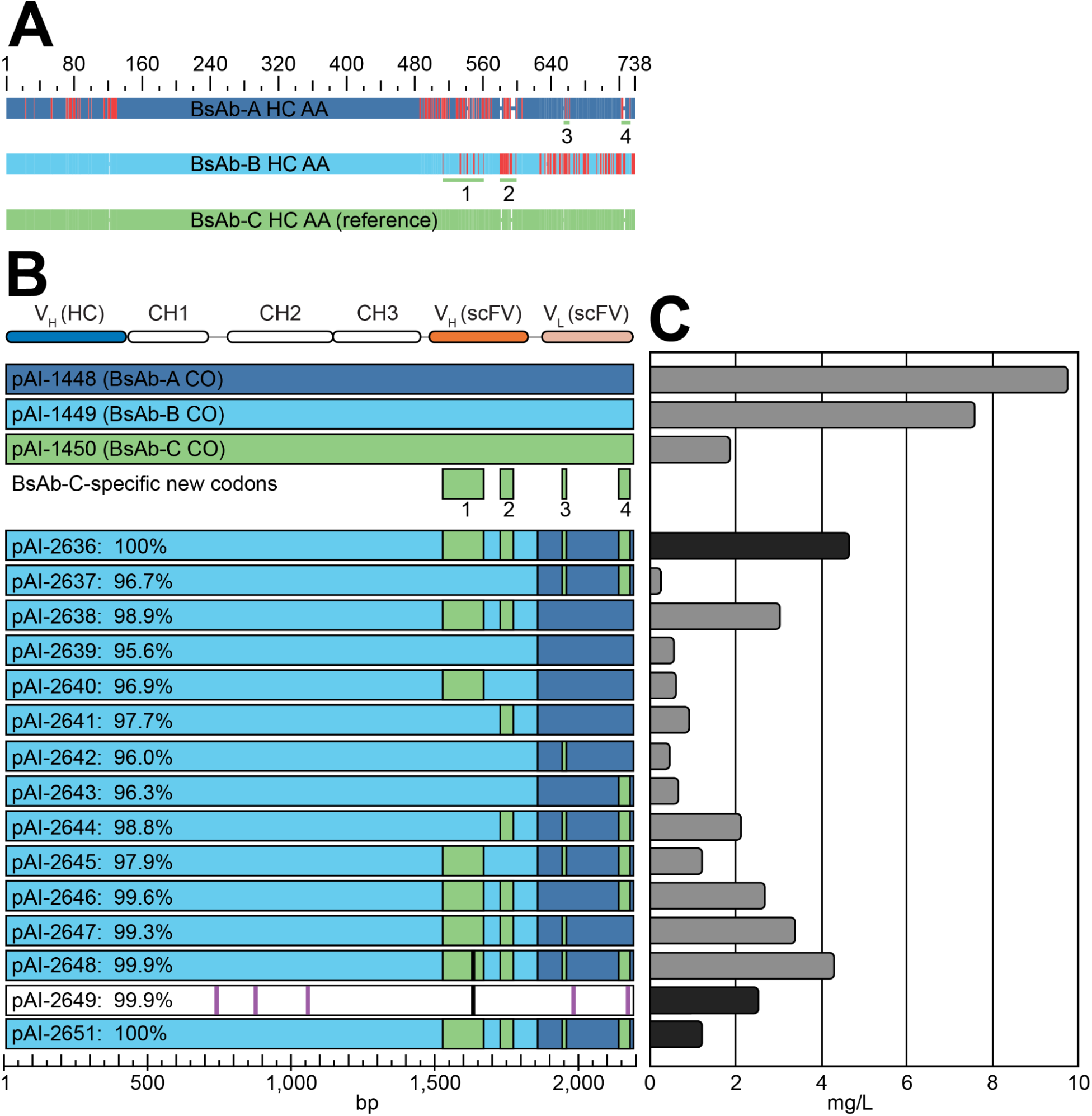
bsAb transient production with revised bsAb-C heavy chain designs A) MAFFT alignment of amino acids sequences for bsAb-A HC, bsAb-B HC, and bsAb-C HC, with bsAb-C as the reference. Blue and cyan indicate where bsAb-A and bsAb-B amino acids match bsAb-C, while red indicates amino acid mismatches or indels relative to bsAb-C.Four nucleotide stretches where the bsAb-C HC coding sequence is unique compared to the other bsAbs are shown as numbered green lines. B) Visualization of revised bsAb-C HC DNA sequences and sequence variants. DNA sequences for bsAb-A HC (pAI-1448; blue) and bsAb-B HC (pAI-1449; cyan) were used as the basis for bsAb-C HC DNA sequences or sequence variants, because the amino acid sequence for bsAb-C HC is almost identical to bsAb-B HC in the N-terminal portion and almost identical to bsAb-A HC in the C-terminal portion. Four stretches of DNA sequences specific to bsAb-C HC are shown as numbered green boxes. Green stretches use codons derived from bsAb-A HC CO (stretches 3 and 4) or bsAb-B HC CO (stretches 1 and 2) for matching amino acids and bsAb-C HC CO for amino acids that are unique to bsAb-C HC. Sequences are either identical to bsAb-C HC at the amino acid level (pAI-2636 and pAI-2651) or chimeric, being mostly identical to bsAb-C HC but with some stretches of bsAb-C HC-specific amino acid sequence being replaced with bsAb-A HC or bsAb-B HC sequence. Percentages refer to the amino acid identity of the variant compared to the reference bsAb-C HC. Thick purple vertical lines (pAI-2649) denote additional changes to splicing motifs. Thick black vertical lines (pAI-2648 and pAI-2649) denote amino acid change from RDK to RDN. C) Titers for bsAb-C HC variants. Titers were measured after 3 days of culture following transient transfection. Darker bars indicate data points measured in a different experiment and normalized to grey bar titers based on four samples shared between the experiments. The bsAb-C sequence variants were combined with bsAb-B LC due to identical amino acid sequence for both light chains.

After screening the panel of bsAb-C variants transiently, we proceeded to stable integration of a subset of four variants encompassing all combinations of original and chimeric codon usage and RDK/RDN amino acid motifs. Furthermore, to isolate the effects of codon usage and ribosomal pausing from other variables such as light chain codon usage, two new constructs (pAI-3656 and pAI-3657) were generated. These use the bsAb-C heavy chain with the original codon optimization from pAI-1450, paired with the bsAb-B light chain, which is amino acid identical to the bsAb-C light chain and matches the expression format of pAI-2636 and pAI-2648. pAI-3656 and pAI-3657 differ only by inclusion of the RDK or RDN motif, respectively. Stable lines were created by random integration followed by selection in glutamine-free medium. After stable integration and 6 day batch production culture, we observed titer increases of 1.9x and 4.4x with RDK to RDN mutation and new codons and respectively, compared to the original bsAb-C sequence. Combining both new codons and the RDK to RDN change (pAI-2648) boosted titer by ∼11x compared to the original bsAb-C (pAI-3656). Similar trends were observed with a fed batch production run (Supplementary Figure S9).

**Figure 5:**
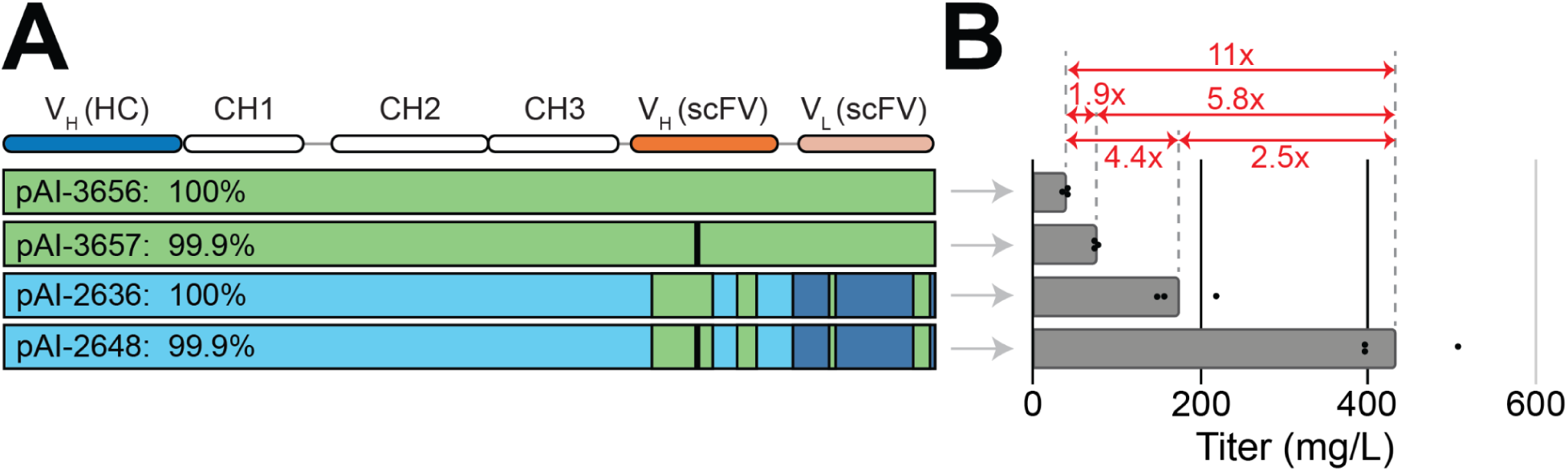
Multiplicative titer improvements in stable bispecific production driven by RDK to RDN amino acid change or chimeric codons. A) HC sequence comparison. Percentages indicate amino acid identity to bsAb-C HC. pAI-3656 is bsAb-C HC with the original RDK motif. pAI-3657 is bsAb-C HC with RDN (denoted by the black vertical line). pAI-2636 is bsAb-C HC with chimeric codons. pAI-2648 is bsAb-C HC with both the RDK to RDN change and chimeric codons. The LC nucleotide sequence is identical for all four expression plasmids (schematic not shown). B) Octet titers from 6 day stable pool batch production cultures. Three replicates are indicated as dots and the average is plotted as bars. The RDK to RDN and the chimeric codon changes result in a combined 11x improvement in bsAb-C titer, with RDK to RDN accounting for between 1.9x-2.5x and codon changes accounting for between 4.4x-5.8x of the increase.

## Discussion

In this study, we diagnosed the root cause for low-production of a bispecific antibody, and then applied these findings to improve productivity. Our results point to two significant contributors to low titers: codon usage within the heavy chain scFv despite undergoing standard codon optimization and ribosomal pausing at a specific position in the heavy chain. Resolving these two factors resulted in a multiplicative ∼11-fold increase in titer, increasing expression of the full-length bsAb to levels comparable to those observed for two other bispecifics with similar IgG-scFv architectures and with sufficient productivity to make the molecule viable for further development.

The data suggest additional considerations and recommendations for bispecific antibody design. Specifically, running different codon optimized variants in parallel where possible, ideally with different optimizers or optimization parameters, can mitigate the chance that problematic codon choice will impact all sequences for a given bispecific. We also recommend screening coding sequences for R-X-K class ribosomal pause sites and removing them if possible. Reducing the occurrence of P-P/G/D ribosomal pause sites could be considered but might not be feasible due to how commonly they can occur.

These findings have broad implications for both the design and optimization of recombinant antibody therapeutics, particularly in early-stage development pipelines where molecule variants are often quickly eliminated. By investigating the contributions of nucleotide and amino-acid level features to protein expression during antibody engineering, it is possible to more effectively balance functional design with manufacturability, leading to the potential rescue of otherwise promising candidates that might be eliminated due to suboptimal expression and enabling more efficient progression through the pipeline. Importantly, this study highlights how relatively modest sequence changes such as codon optimization and targeted motif edits can dramatically shift expression potential, potentially providing a workflow for troubleshooting other low-expressing constructs.

Through this study we also demonstrate the utility of a bioinformatics-based approach to troubleshooting production issues with biologics. Using RNA-seq, structure predictions, and sequence level analysis, numerous hypotheses for low production were provisionally ruled out, while others were prioritized for further investigation in the lab. Specifically, the workflow applied here included analysis of differential gene and gene set expression, splicing isoforms, transcriptional resource allocation, miRNA target sites, codon choice and optimization, ribosome collision motifs, and RNA structure. This integrative process enabled the identification of sequence-level adjustments, while deprioritizing factors such as cellular bottlenecks, transcript stability, and RNA structure.

Looking forward, integrating predictive tools with early expression testing may transform how we optimize antibody design and construct generation. Sequence-aware strategies supported by codon usage data, ribosomal profiling, and translation modeling can identify expression liabilities earlier. Pairing this approach with rapid screening of variants earlier in development may improve manufacturability while preserving function, especially for complex constructs like bispecifics. As the field increasingly shifts toward multi-specific antibody formats, these methods will become essential for avoiding delays later in development. Combining computational predictions with rational, small-scale variant testing offers a scalable path to ensure both function and expressibility are maintained from the outset.

## Acknowledgements

We thank Rene Hubert and Eric Gislason at Amgen for their insights on CHO cell lines and molecules selected for use in the transcriptomics studies.

## Supplementary Figures and Captions

**Supplementary Figure S1:**
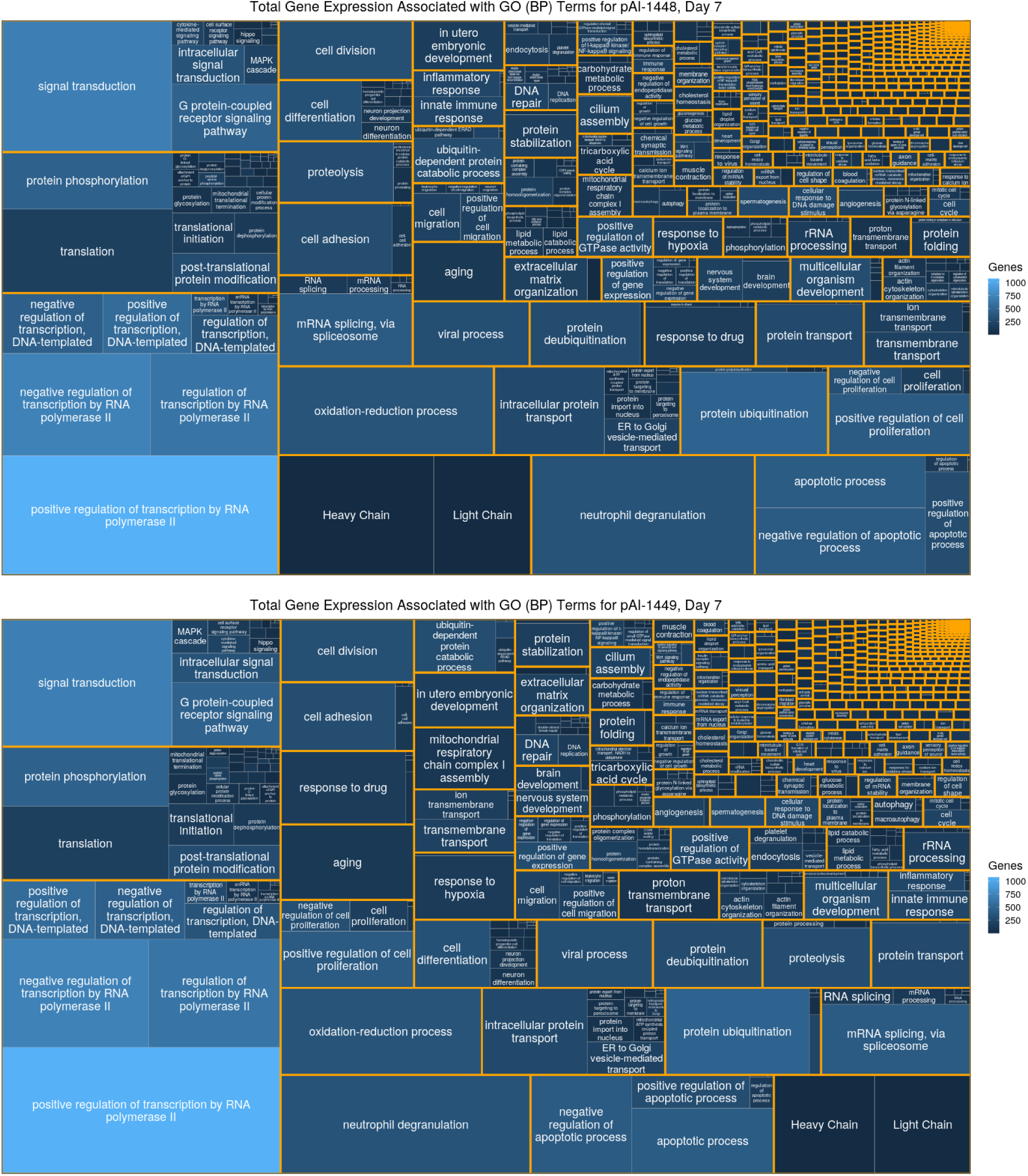
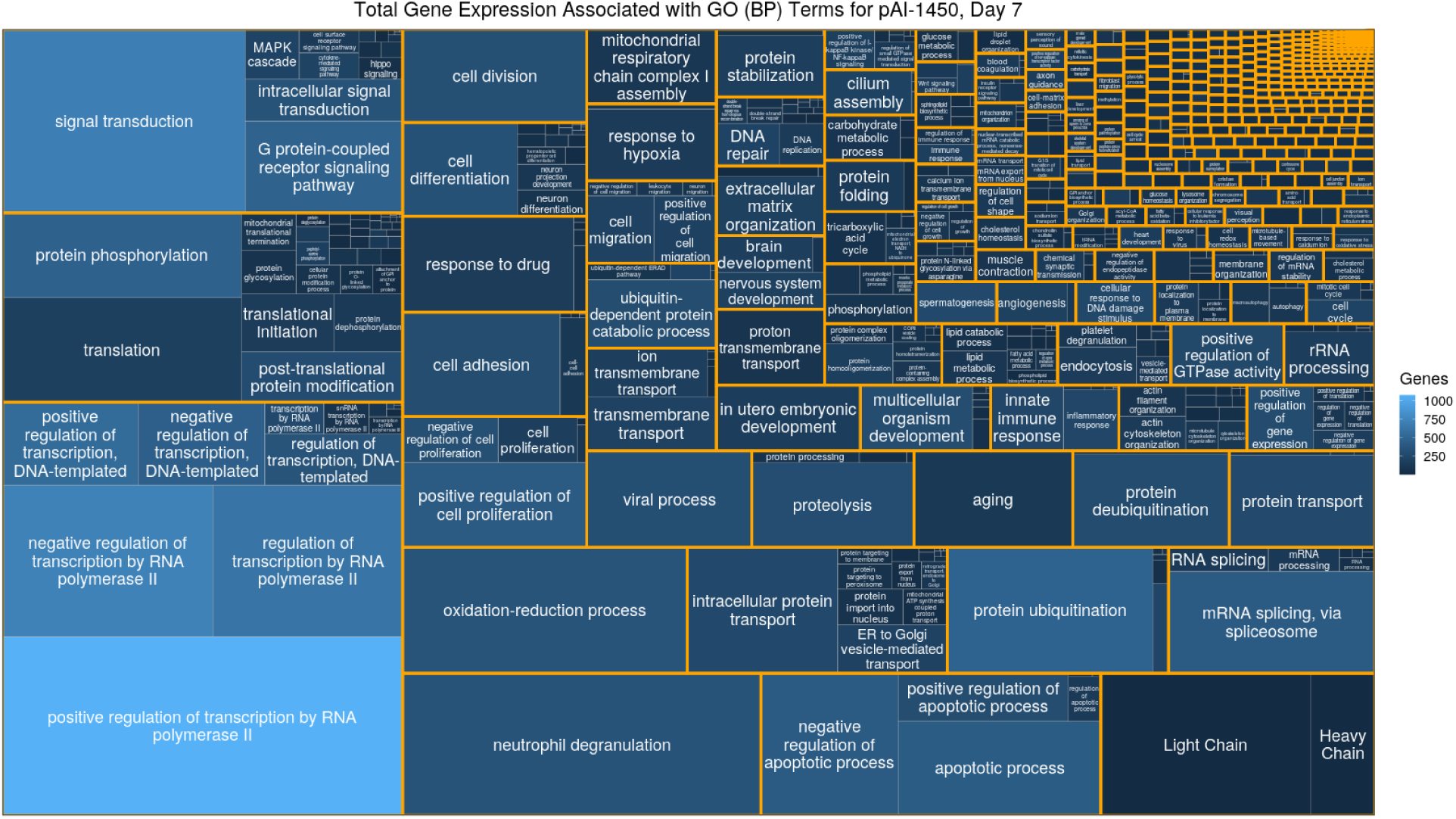
Treemap of gene expression by GO terms We generated treemaps to visualize the resources allocated to express genes associated with different GO-terms for the three bispecific molecules at day 7. [35] bsAb-A (pAI-1448) is shown on the first, bsAb-B (pAI-1449) is shown second, and bsAb-C (pAI-1450) is shown third. The size of the rectangles represents total FPKM of genes associated with the GO-terms (shown as labels). The color of the rectangles corresponds to the number of genes associated with each GO-term. Overall gene expression for different GO terms is similar across the three bispecifics, with the major discrepancy between the amount of HC and LC.

**Supplementary Figure S2:**
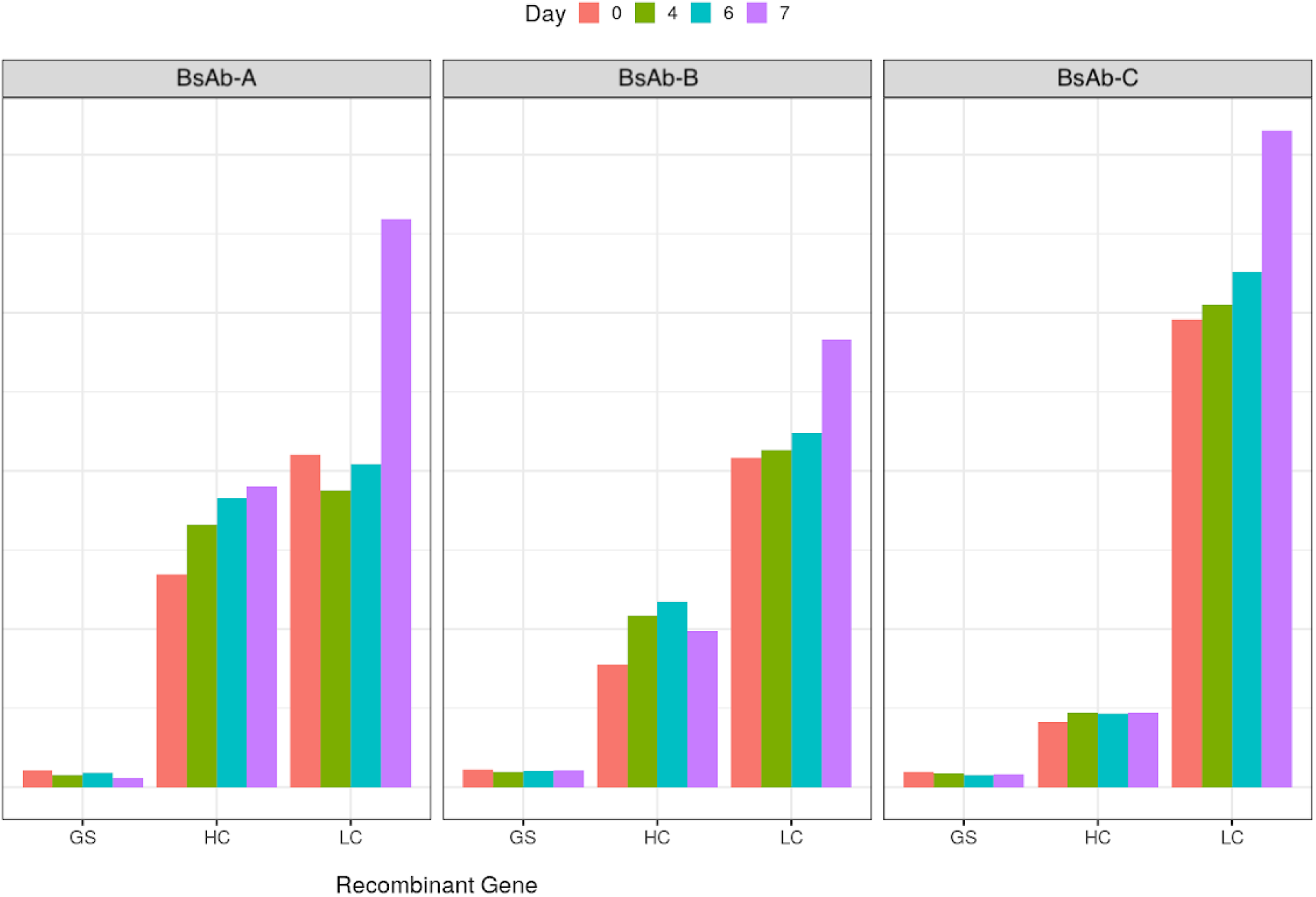
Gene expression analysis of bispecific expression construct genes Transcript per million (TPM) values were calculated for glutamine synthetase (GS), heavy chain (HC), and light chain (LC) across the three different bispecifics on four different days. LC expression was higher than HC for all bispecifics. GS expression was low and consistent, due to the use of a weaker promoter. HC expression was especially low for bsAb-C HC at less than 5000 TPM.

**Supplementary Figure S3:**
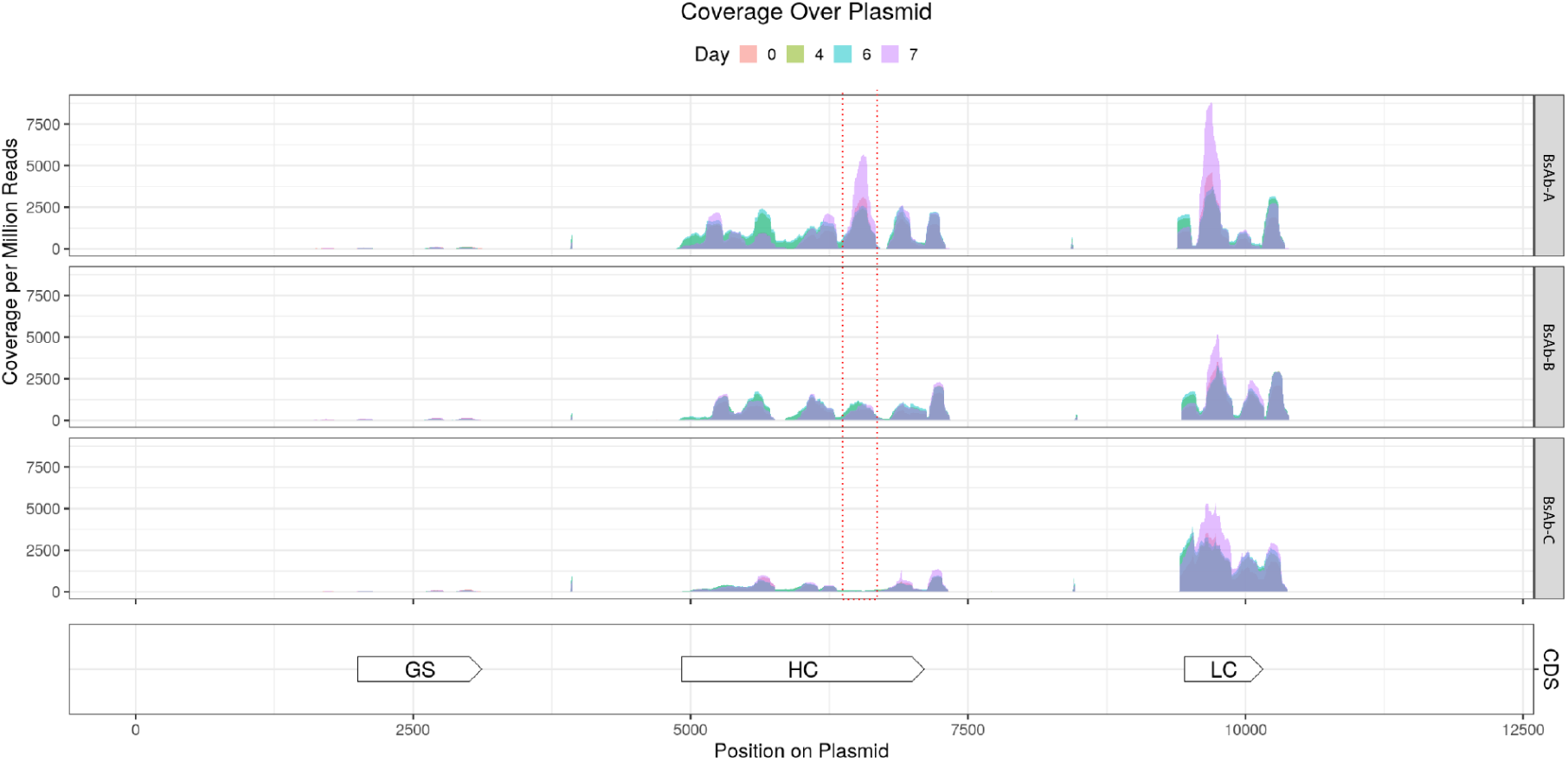
Read coverage plots for the bispecific expression constructs Read coverage per million reads is plotted across all positions in the three bispecific plasmids across four days. GS, HC, and LC positions within the plasmids are denoted in the bottom graph. A region with low coverage in bsAb-C HC relative to the other bispecifics is demarcated by red dashed lines.

**Supplementary Figure S4.A:**
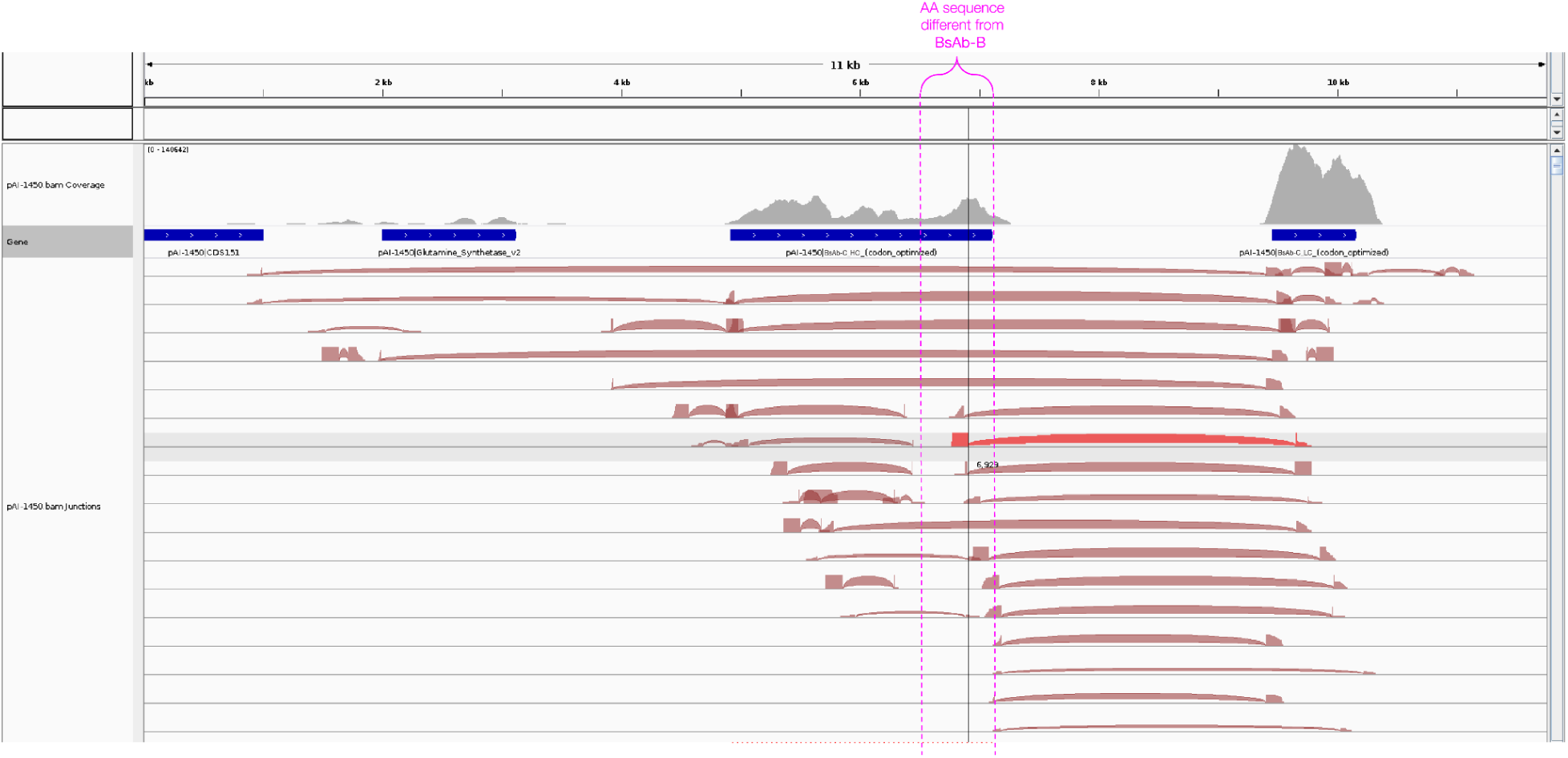
Visualization of RNA splicing junction reads BAM files were loaded into Integrative Genomics Viewer (IGV) and junction reads were visualized. Note that many of the junction reads shown in IGV are mapping artifacts due to repeated regions within the plasmid sequence. To avoid false positives due to mapping, junction reads were filtered by requiring that they do not overlap repeated regions and visualized using ggplot2.

**Supplementary Figure S4.B:**
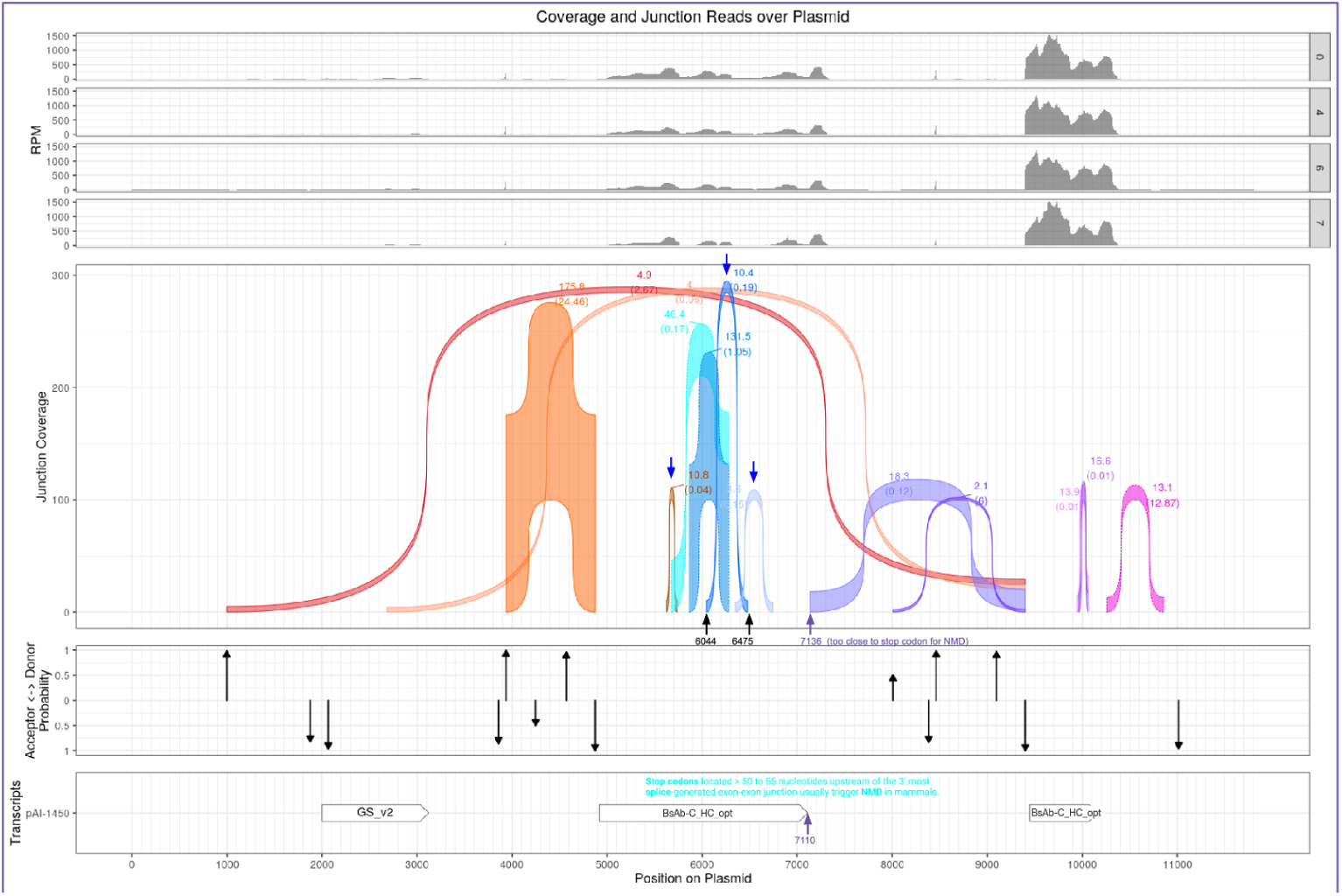
Visualization of RNA splicing junction reads continued Visualizations of splicing within the bsAb-C expression plasmid, with ribbons representing filtered junction reads and arrows representing spliceAI-predicted splice site probabilities.

**Supplementary Figure S5:**
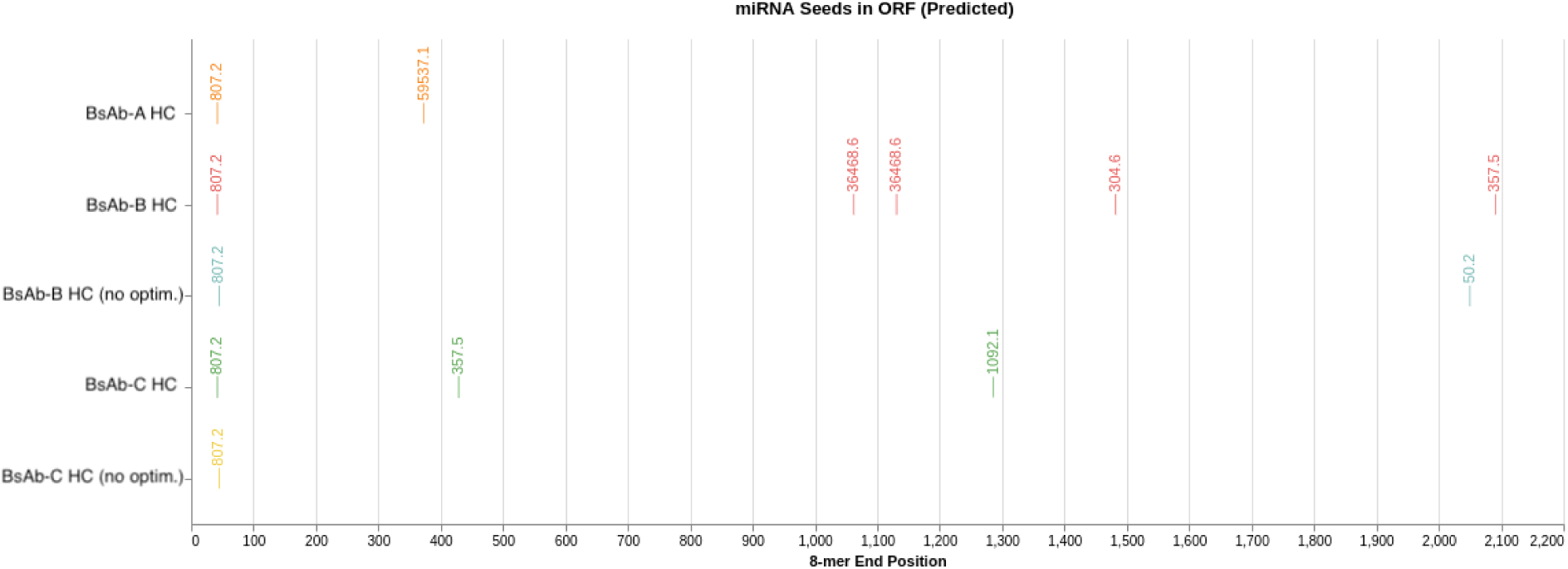
miRNA RPMs We plotted the combined RPMs of microRNAs sharing the same seed sequence expressed in CHO. The RPMs were from a public data set. [36]

**Supplementary Figure S6:**
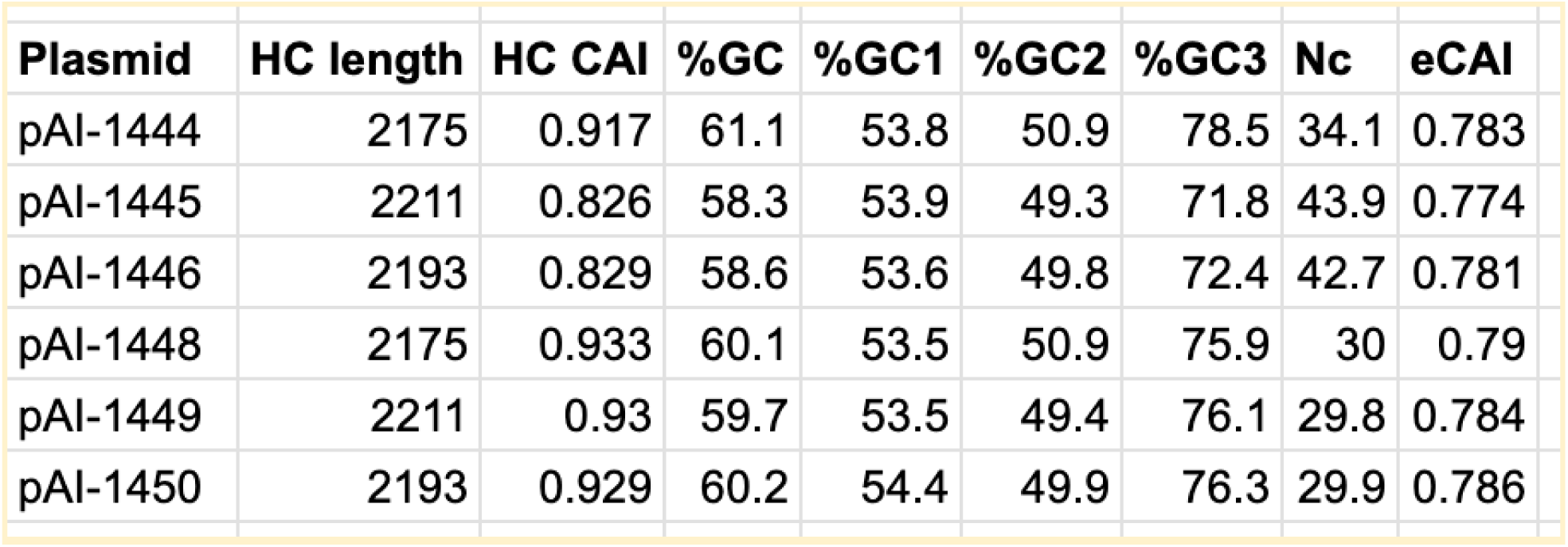
Codon usage statistics CAI, %GC, %GC at the first position in each codon, %GC at the second position in each codon, %GC at the third position in each codon, number of effective codons, and effective CAI were calculated for each heavy chain sequence using the codon usage table for Cricetulus griseus. [37] Calculations were made using CAIcal. [38]

**Supplementary Figure S7:**
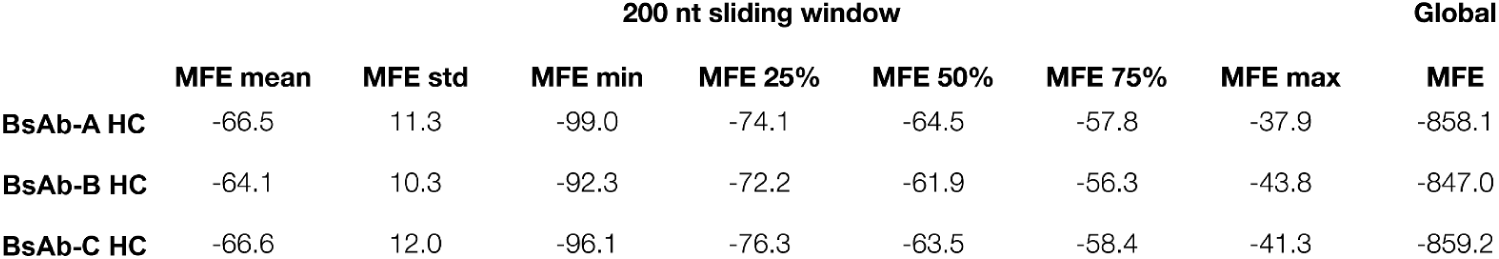
RNA folding statistics Statistics of MFE of sequences in a 200-nt sliding window of the mRNA or MFE of the full mRNA sequence (last column). The MFE was calculated using the RNAfold function of the ViennaRNA package.

**Supplementary Figure S8:**
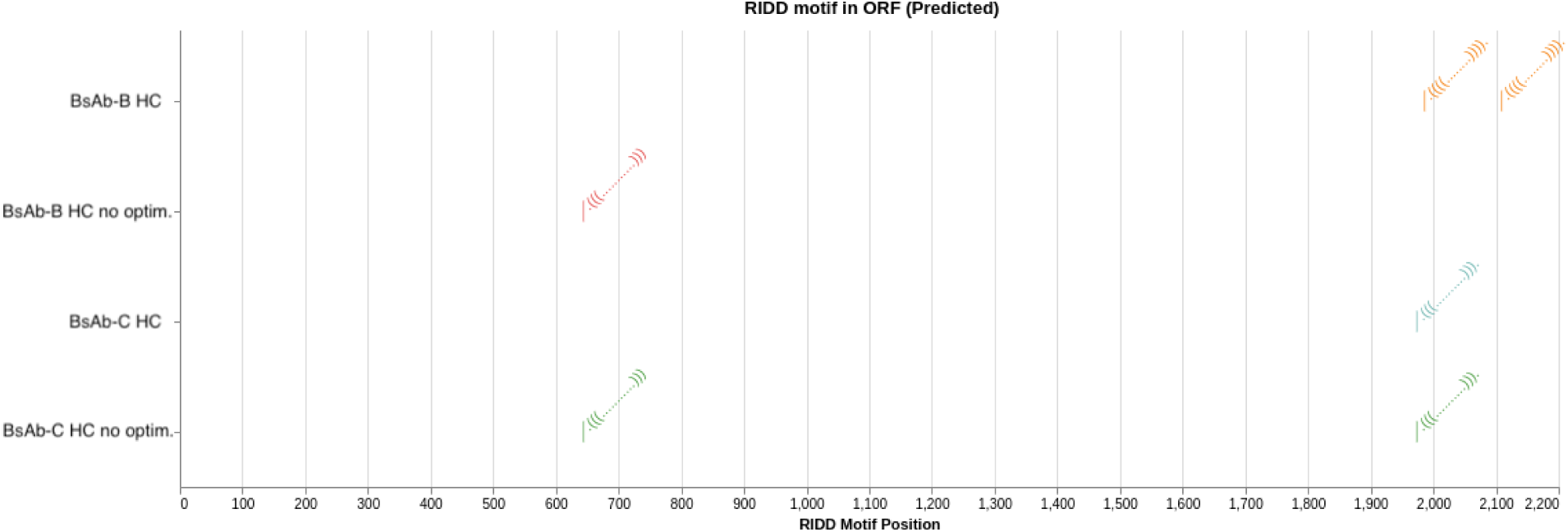
Predicted stem-loop endomotifs for RIDD. The Stem-loop endomotifs were identified by analyzing RNA structures predicted using the RNAfold function of the ViennaRNA package. [39]

**Supplementary Figure S9:**
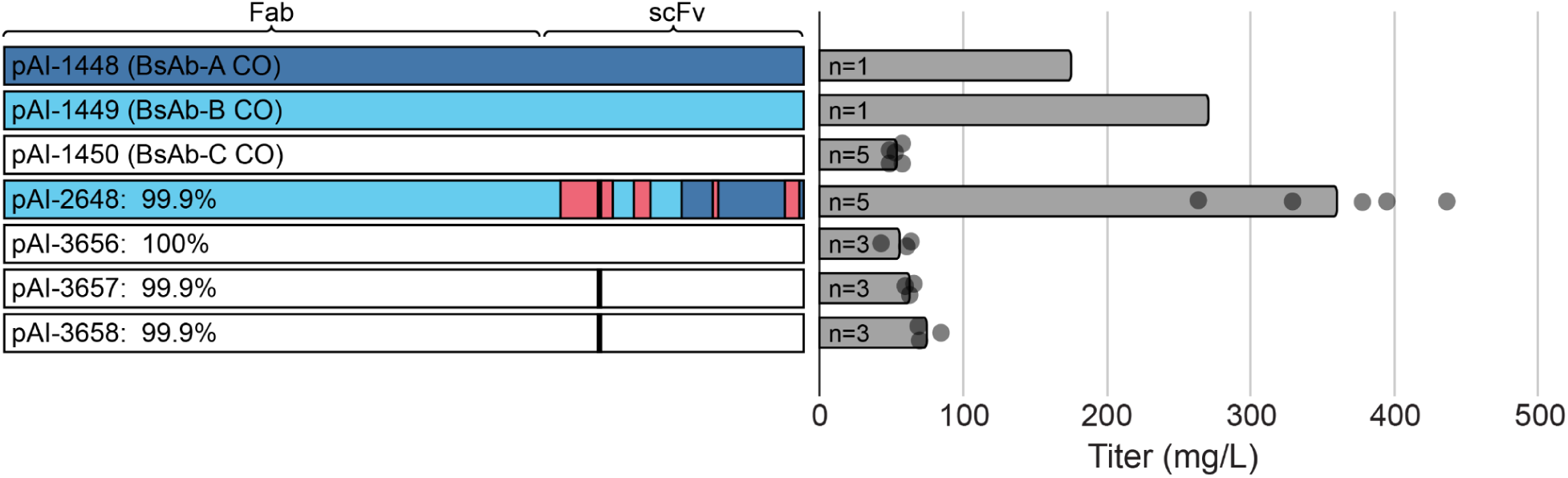
Fed batch shake flask titer confirmation driven by RDK to RDN amino acid change or new codons. A) Heavy chain sequence comparison. Percentages indicate amino acid identity to bsAb-C HC. pAI-1448 is bsAb-A codon optimized. pAI-1449 is bsAb-B codon optimized. pAI-1450 is bsAb-C codon optimized. pAI-2648 is bsAb-C HC with RDK to RDN change and new codons. pAI-3656 is bsAb-C HC with RDK. pAI-3657 is bsAb-C HC with RDN. pAI-3658 is bsAb-C HC with RDN Version 2. B) Octet titers from 14 day fed batch cultures. N denotes replicates for each plasmid while replicates are indicated as dots with the average plotted as bars. The RDK to RDN and the new codon changes result in a combined 11x improvement in bsAb-C titer, with RDK to RDN accounting for between 1.9x-2.5x and codon changes accounting for between 4.4x-5.8x of the increase.

## References

1. Lyu, X., Zhao, Q., Hui, J., Wang, T., Lin, M., Wang, K., Zhang, J., Shentu, J., Dalby, P. A., Zhang, H., & Liu, B. (2022). The global landscape of approved antibody therapies. Antibody therapeutics, 5(4), 233–257. 10.1093/abt/tbac021

2. Chae, Y. K., Arya, A., Iams, W., Cruz, M. R., Chandra, S., Choi, J., & Giles, F. (2018). Current landscape and future of dual anti-CTLA4 and PD-1/PD-L1 blockade immunotherapy in cancer; lessons learned from clinical trials with melanoma and non-small cell lung cancer (NSCLC). Journal for immunotherapy of cancer, 6(1), 39. 10.1186/s40425-018-0349-3

3. Ma, J., Mo, Y., Tang, M., Shen, J., Qi, Y., Zhao, W., Huang, Y., Xu, Y., & Qian, C. (2021). Bispecific Antibodies: From Research to Clinical Application. Frontiers in immunology, 12, 626616. 10.3389/fimmu.2021.626616

4. Wei, J., Yang, Y., Wang, G., & Liu, M. (2022). Current landscape and future directions of bispecific antibodies in cancer immunotherapy. Frontiers in immunology, 13, 1035276. 10.3389/fimmu.2022.1035276

5. KuicK Research. (2025, April 4). Bispecific trispecific antibodies market size, FDA approval, clinical trials & drug sales insight 2030 [Press release]. Retrieved from https://www.globenewswire.com/news-release/2025/04/04/3056061/0/en/Bispecific-Trispecific-Antibodies-Market-Size-FDA-Approval-Clinical-Trials-Drug-Sales-Insight 2030.html

6. Jayapal, Karthik & Wlaschin, K.F. & Hu, W.S. & Yap, M.G.S.. (2007). Recombinant Protein Therapeutics from CHO Cells - 20 Years and Counting. Chemical Engineering Progress. 103. 40–47.

7. Ding, M., Shen, L., Xiao, L., Liu, X., & Hu, J. (2021). A cell line development strategy to improve a bispecific antibody expression purity in CHO cells. Biochemical Engineering Journal, 166, 107857. 10.1016/j.bej.2020.107857

8. Donaldson, J., Kleinjan, D. J., & Rosser, S. (2022). Synthetic biology approaches for dynamic CHO cell engineering. Current opinion in biotechnology, 78, 102806. 10.1016/j.copbio.2022.102806

9. Zeh, N., Bräuer, M., Raab, N., Handrick, R., & Otte, K. (2022). Exploring synthetic biology for the development of a sensor cell line for automated bioprocess control. Scientific reports, 12(1), 2268. 10.1038/s41598-022-06272-x

10. Kumar, D., Gangwar, N., Rathore, A. S., & Ramteke, M. (2022). Multi-objective optimization of monoclonal antibody production in bioreactor. Chemical Engineering and Processing - Process Intensification, 180, 108720. 10.1016/j.cep.2021.108720

11. Rader, C. (2011). DARTs take aim at BiTEs. Blood, 117(17), 4403–4404. 10.1182/blood-2011-02-337691

12. Wang, Q., Chen, Y., Park, J., Liu, X., Hu, Y., Wang, T., McFarland, K., & Betenbaugh, M. J. (2019). Design and Production of Bispecific Antibodies. Antibodies (Basel, Switzerland), 8(3), 43. 10.3390/antib8030043

13. Magistrelli, G., Poitevin, Y., Schlosser, F., Pontini, G., Malinge, P., Josserand, S., Corbier, M., & Fischer, N. (2017). Optimizing assembly and production of native bispecific antibodies by codon de-optimization. mAbs, 9(2), 231–239. 10.1080/19420862.2016.1267088

14. Zhang, W., Wang, H., Feng, N., Li, Y., Gu, J., & Wang, Z. (2022). Developability assessment at early-stage discovery to enable development of antibody-derived therapeutics. Antibody Therapeutics, 6(1), 13–29. 10.1093/abt/tbac029

15. Fawcett, C., Tickle, J. R., & Coles, C. H. (2024). Facilitating high throughput bispecific antibody production and potential applications within biopharmaceutical discovery workflows. mAbs, 16(1), 2311992. 10.1080/19420862.2024.2311992

16. Brinkmann, U., & Kontermann, R. E. (2017). The making of bispecific antibodies. mAbs, 9(2), 182–212. 10.1080/19420862.2016.1268307

17. Labrijn, A.F., Janmaat, M.L., Reichert, J.M. et al. Bispecific antibodies: a mechanistic review of the pipeline. Nat Rev Drug Discov 18, 585–608 (2019). 10.1038/s41573-019-0028-1

18. Merchant, A., Zhu, Z., Yuan, J. et al. An efficient route to human bispecific IgG. Nat Biotechnol 16, 677–681 (1998). 10.1038/nbt0798-677

19. Gunasekaran, K., Pentony, M., Shen, M., Garrett, L., Forte, C., Woodward, A., Ng, S. B., Born, T., Retter, M., Manchulenko, K., Sweet, H., Foltz, I. N., Wittekind, M., & Yan, W. (2010). Enhancing antibody Fc heterodimer formation through electrostatic steering effects: Applications to bispecific molecules and monovalent IgG. Journal of Biological Chemistry, 285(25), 19637–19646. 10.1074/jbc.M110.117382

20. Klein, C., Schaefer, W., & Regula, J. T. (2016). The use of CrossMAb technology for the generation of bi- and multispecific antibodies. mAbs, 8(6), 1010–1020. 10.1080/19420862.2016.1197457

21. Ong, H. K., Nguyen, N. T. B., Bi, J., & Yang, Y. (2022). Vector design for enhancing expression level and assembly of knob-into-hole based FabscFv-Fc bispecific antibodies in CHO cells. Antibody therapeutics, 5(4), 288–300. 10.1093/abt/tbac025

22. Yoon, C., Lee, E. J., Kim, D., Joung, S., Kim, Y., Jung, H., Kim, Y. G., & Lee, G. M. (2024). SiMPl-GS: Advancing Cell Line Development via Synthetic Selection Marker for Next-Generation Biopharmaceutical Production. Advanced science (Weinheim, Baden-Wurttemberg, Germany), 11(38), e2405593. 10.1002/advs.202405593

23. Zhang, J., Cao, W., Yu, L., Cui, Y., Xu, K., Tian, J., Hogl, S., Kaufmann, H., Zhou, W., & Gu, S. (2024). Stepwise cell culture process intensification for high-productivity and cost-effective commercial manufacturing of a Mabcalin™ bispecifics. Biochemical Engineering Journal, 211, 109476. 10.1016/j.bej.2024.109476

24. Jansen, R., Bussemaker, H. J., & Gerstein, M. (2003). Revisiting the codon adaptation index from a whole-genome perspective: Analyzing the relationship between gene expression and codon occurrence in yeast using a variety of models. Nucleic Acids Research, 31(8), 2242–2251. 10.1093/nar/gkg306

25. Lorenz, R., Bernhart, S.H., Höner zu Siederdissen, C., et al. ViennaRNA Package 2.0. Algorithms Mol Biol 6, 26 (2011). 10.1186/1748-7188-6-26

26. Le Thomas, A., Ferri, E., Marsters, S. et al. Decoding non-canonical mRNA decay by the endoplasmic-reticulum stress sensor IRE1α. Nat Commun 12, 7310 (2021). 10.1038/s41467-021-27597-7

27. Kontermann, R. E., & Brinkmann, U. (2015). Bispecific antibodies. Drug Discovery Today, 20(7), 838–847. 10.1016/j.drudis.2015.02.008

28. Coloma, M. J., & Morrison, S. L. (1997). Design and production of novel tetravalent bispecific antibodies. Nature biotechnology, 15(2), 159–163. 10.1038/nbt0297-159

29. Zuallaert, J., Godin, F., Kim, M., Soete, A., Saeys, Y., & De Neve, W. (2018). SpliceRover: Interpretable convolutional neural networks for improved splice site prediction. Bioinformatics, 34(24), 4180–4188. 10.1093/bioinformatics/bty497

30. Zhao, T., Chen, Y. M., Li, Y., et al. (2021). Disome-seq reveals widespread ribosome collisions that promote cotranslational protein folding. Genome Biology, 22, 16. 10.1186/s13059-020-02256-0

31. Simms, C. L., Yan, L. L., Qiu, J. K., & Zaher, H. S. (2019). Ribosome Collisions Result in +1 Frameshifting in the Absence of No-Go Decay. Cell reports, 28(7), 1679–1689.e4. 10.1016/j.celrep.2019.07.046

32. Juszkiewicz, S., Slodkowicz, G., Lin, Z., Freire-Pritchett, P., Peak-Chew, S. Y., & Hegde, R. S. (2020). Ribosome collisions trigger cis-acting feedback inhibition of translation initiation. eLife, 9, e60038. 10.7554/eLife.60038

33. Han, P., Shichino, Y., Schneider-Poetsch, T., Mito, M., Hashimoto, S., Udagawa, T., Kohno, K., Yoshida, M., Mishima, Y., Inada, T., & Iwasaki, S. (2020). Genome-wide Survey of Ribosome Collision. Cell reports, 31(5), 107610. 10.1016/j.celrep.2020.107610

34. Barnes, L. M., Bentley, C. M., & Dickson, A. J. (2003). Stability of protein production from recombinant mammalian cells. Biotechnology and Bioengineering, 81, 631–639. 10.1002/bit.10517

35. R Graph Gallery. (n.d.). Treemap [Web page]. From https://r-graph-gallery.com/treemap.html

36. Hammond, S., Swanberg, J. C., Polson, S. W., & Lee, K. H. (2012). Profiling conserved microRNA expression in recombinant CHO cell lines using Illumina sequencing. Biotechnology and bioengineering, 109(6), 1371–1375. 10.1002/bit.24415

37. Kazusa Codon Usage Database. (n.d.). Codon usage table — Species code 10029. Retried from https://www.kazusa.or.jp/codon/cgi-bin/showcodon.cgi?species=10029

38. Puigbò, P., Bravo, I.G. & Garcia-Vallve, S. CAIcal: A combined set of tools to assess codon usage adaptation. Biol Direct 3, 38 (2008). 10.1186/1745-6150-3-38

39. Le Thomas, A., Ferri, E., Marsters, S., Harnoss, J. M., Lawrence, D. A., Zuazo-Gaztelu, I., Modrusan, Z., Chan, S., Solon, M., Chalouni, C., Li, W., Koeppen, H., Rudolph, J., Wang, W., Wu, T. D., Walter, P., & Ashkenazi, A. (2021). Decoding non-canonical mRNA decay by the endoplasmic-reticulum stress sensor IRE1α. Nature communications, 12(1), 73 10.10.1038/s41467-021-27597-7

